# *Aedes* mosquito distribution across urban and peri-urban areas of Kinshasa city, Democratic Republic of Congo

**DOI:** 10.1101/2025.09.03.674006

**Authors:** Victoire Nsabatien, Josue Zanga, Nono Mvuama, Arsene Bokulu, Hyacinthe Lukoki, Glodie Diza, Dorcas Kantin, Leon Mbashi, Christelle Bosulu, Narcisse Basosila, Erick Bukaka, Fiacre Agossa, Jonas Nagahuedi, Jean Claude Palata, Emery Metelo

**Author notes:** **Corresponding author(s):** Victoire Nsabatien.

## Abstract

In the Democratic Republic of Congo (DRC) *Aedes* mosquitoes are principal vectors of medically important arboviruses, with major implications for yellow fever, chikungunya and dengue. However, systematic surveillance of these species remains limited, constrained by competing public health priorities such as malaria and other neglected tropical diseases. This gap in surveillance prevents the rapid detection of changes in the distribution, abundance and behaviour, particularly in rapidly urbanizing environments where breeding habitats are proliferating and ecological conditions are favourable for the establishment of these vectors. To address this gap, spatially explicit, small-scale data on *Aedes* populations in urban and peri-urban areas are needed to accurately assess transmission risk and develop targeted, evidence-based vector control strategies. Here, we present a geo-referenced dataset of 6,577 entomological occurrence records collected in 20224 throughout urban and peri-urban areas of Kinshasa city, DRC, using Larval dipping, Human landing catches, Prokopack aspirator, and BG-Sentinel traps. Records include *Aedes albopictus* (n = 2,694), *Aedes aegypti* (n = 1939), *Aedes vittatus* (n = 2), and *Aedes* spp. (n = 1,942), each annotated with species, sex, life stage, reproductive status, and spatial coordinates. The dataset is published as a Darwin Core archive in the Global Biodiversity Information Facility (GBIF), and represents the most detailed, spatially explicit record of *Aedes* mosquito occurrence in Kinshasa to data, providing a robust foundation for entomological and modelling research to support data driven arbovirus vector control strategies in DRC.

**Subject Areas:** Ecology, Biodiversity, Vector Biology

## Data description

## Background and context

The spread of arbovirus vectors such as *Aedes aegypti* (Linnaeus, 1762) (Diptera: Culicidea) and *Aedes albopictu*s (Skuse, 1895) (Diptera: Culicidea) is accelerating across Africa, driven by human mobility, expanding transport networks, urbanisation and climate change [1-4]. These species are now established across Africa countries and have played a major role in the transmission of yellow fever virus, chikungunya virus and dengue virus in Central African countries such as Cameroon, Gabon, the Central African Republic, the Republic of Congo and the Democratic Republic of Congo [5-10].

In the DRC, although no nationwide survey of their distribution has been conducted, limited entomological studies and global distribution models based on environmental variables without entomological data indicate that *Ae. Aegypti* is widespread, whereas *Ae. Albopictus* remains largely restricted to the western regions, where it increasingly displaces *Ae. Aegypti* [11-13]. In Kinshasa City, the introduction in 2018 has led the co-occurrence of both species in urban and peri-urban areas, increasing the risk of arbovirus transmission. *Ae. aegypti* is more common in densely populated urban areas with high building density, where it prefers to reproduce in artificial containers, while *Ae. albopictus* is more frequent in peri-urban and rural areas, where it prefers to reproduce in containers surrounded by vegetation [11-15].

Here we present recent data on the geographical distribution of *Aedes* species and abundance of *Aedes* species across Kinshasa, DRC, collected between January and December 2024.

## Methods

### General spatial coverage

This study was conducted in two areas with contrasting levels of urbanisation in Kinshasa city. Mont-Ngafula, a peri urban area in the south-west of the city located between latitude 4°15’ S and longitude 15°14’ E; and Kitambo, an urban area in the north-west of the city located between latitude 04°20’S and longitude 15°16′ E. Within each area, two sampling sites were selected.

**Figure 2.**
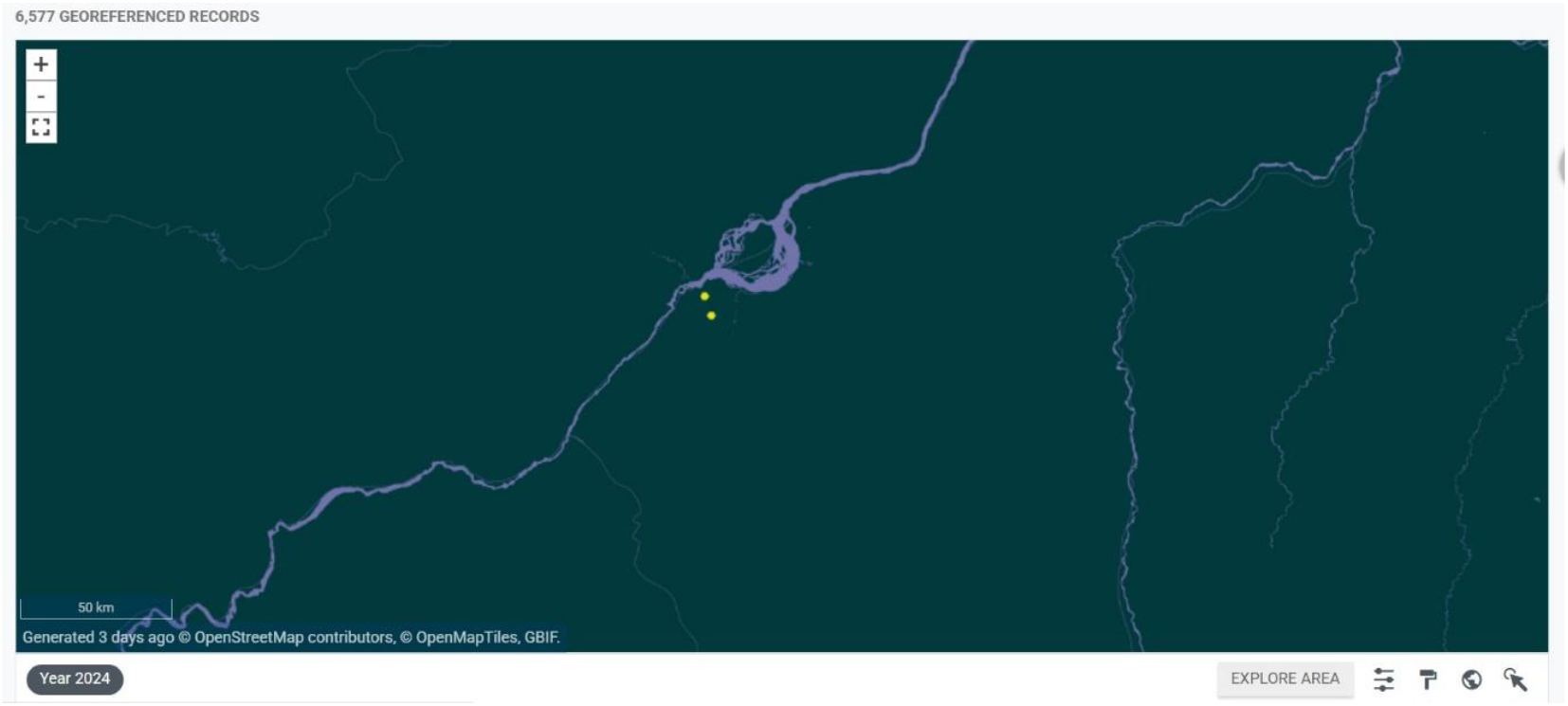
Interactive map of the georeferenced occurrences hosted by GBIF [16].

### Mosquito collection

The general taxonomic coverage description for this work is the Culicidae Family, specifically *Ae. aegypti* (commonly known as yellow fever mosquito; NCBI:txid7159*), Ae. albopictus* (commonly known as the Asian tiger mosquito or moustique tigre in French; NCBI:txid7160) and *Ae. vittatus* (Bigot, 1861) (formerly known as *Culex vittatus*; NCBI:txid317808), and *Aedes* spp., where larval specimens were performed only to the genus level.

Four sampling techniques were used to collect immature and adult stages of *Aedes* between January 11, 2024 - December 20, 2024, covering both dry and rainy seasons.

Immature mosquito stages were collected from potential breeding sites and identified to the genus level, whereas adults collected from different households for each of the three samplings techniques (Human landing catches, Prokopack aspirator and BG-Sentinel trap) were morphologically identified to the species level using the taxonomic keys [17].

### Larval collection

From January to December 2024 study period, immature stages of *Aedes* spp. Were sampled from domestic, peridomestic, and natural habitats using dipping technique once a month. Water from each breeding sites was using with a ladle or a small container to search for Culicidae larvae, specifically *Aedes* spp. Larvae were collected, transferred and stored into jars containing water from their respective breeding sites and transported to the insectary of the Laboratory of Bioecology and Vector Control (BIOLAV), where they were reared to adulthood under insectary conditions (temperature: 28 ± 1°C; relative humidity: 70-80%; photoperiod 14 h: 10 h light: dark photoperiod).

### Human Landing Catches

Human landing catches (HLC) are a widely used method for directly quantifying human-mosquito contact in entomological surveillance [18]. In this study, Adult *Aedes* spp were collected by HLC at each sampling site, with sessions of mosquitoes capture conducted both indoors and outdoors with two groups of collectors, in two periods of times (6:00 a.m. to 12:00 p.m. and 12:00 p.m. to 6:00 p.m). At each collection point, a bare-legged, barefoot volunteer served as bait, collecting mosquitoes using hemolysis tubes. Mosquito samples were then transported to the morphological identification unit of the BIOLAV.

### Prokopack Aspirator

Outdoor-resting *Aedes* mosquitoes were collected using Prokopack aspirator (Model 140, John W.Hock Co., Gainesville, FL, USA) [19]. Every hour, from 6 a.m. to 6 p.m., targeted sampling of potential exophilic resting sites was carried out indoors and outdoors, particularly in crowded areas, under shady vegetation, flowers and aquatic surfaces in aquatic habitats. The collected mosquitoes were placed in small containers labelled by time block and transported to BIOLAV lab.

### BG-Sentinel Trap

Although The BG-Sentinel 2 trap, can operate using mains electricity, the option to run on rechargeable batteries offers a crucial advantage in the regions where access to reliable power is limited. Monthly *Aedes* mosquito collections with the BG-Sentinel 2 (Biogents Mosquito Monitoring), were conducted every hour, from 6 a.m. to 6 p.m, both indoors and outdoors.

### Quality control description

Field work was supervised by a trained entomology technician with one focal point per site to ensure protocol adherence. All equipment was cleaned, inspected, and tested prior to each activity, with battery charged the day before deployment (Prokopack Aspirator and BG-Sentinel 2). After field and laboratory work, and once digitized, the data was validated using the Integrated Publishing Toolkit (IPT) validator tool available from the Global Biodiversity Information Facility (GBIF) [20].

## Results

Overall, 6,577 *Aedes* mosquitoes were collected across all sampling methods (Table 1). *Ae. Albopictus was the most abundant species* (41.0%, n = 2694), followed by *Ae. aegypti* (29.5%, n = 1939), while *Ae. vittatus* was rare (0.03%, n = 2). In addition, 1,942 *Aedes* larvae (29.5%) were collected and not identified to species level. The majority of adult mosquitoes were captured using the prokopack aspirator (n = 2,212) and human landing catches (n = 1,799), with fewer collected by BG-Sentinel traps (624).

**Table 1.** Counts of collected *Aedes* mosquitoes by species across all sampling methods

| Species | Sampling methods |  |  |  |  |
| --- | --- | --- | --- | --- | --- |
|  | Human landing catches<br>n (%) | BG-Sentinel<br>n (%) | Prokopack<br>n (%) | Larva collect<br>n (%) | Total<br>n (%) |
| *Ae. aegypti* | 752 (41.8) | 262 (42.0) | 925 (41.8) | - | 1,939 (29.5) |
| *Ae. albopictus* | 1,046 (58.1) | 362 (58.0) | 1,286 (58.1) | - | 2,694 (41.0) |
| *Ae. vittatus* | 1 (0.1) | - | 1 (0.05) | - | 2 (0.03) |
| *Ae. spp* (*unid) | - | - | - | 1942 (100) | 1,932 (29.5) |
| **Total** | **1,799 (100)** | **624 (100)** | **2,212 (100)** | **1,942 (100)** | **6,577 (100)** |

\*unid : *Aedes* larvae not identified to species level

Species composition showed strong contrasts between urban and peri-urban habitats (Table 2). In Kitambo (urban habitat), *Ae. aegypti* predominated, comprising 40.5% (n = 1,040) of captures during the rainy season and 52.7% (n = 372) in the dry season, whereas *Ae. albopictus* 29.9% (n = 774) and 17.7% (n = 125), respectively. Unidentified *Aedes* spp. contributed nearly one-third of collections in the rainy season (29.5%, n =763). By contrast, Mont-Ngafula (peri-urban habitat) was dominated by *Ae. albopictus*, which accounted for more than half of all specimens in both rainy (53.5%, n = 1,416) and dry (59.3%) seasons.

**Table 2.** Species composition of *Aedes* mosquitoes collected by habitat type and season in Kinshasa.

| Location | Habitat | Season | Species |  |  |  | Total<br>n (%) |
| --- | --- | --- | --- | --- | --- | --- | --- |
|  |  |  | *Ae. aegypti*<br>n (%) | *Ae. albopictus*<br>n (%) | *Ae. vittatus*<br>n (%) | *Ae. spp*<br>n (%) |  |
| Kitambo | Urban | rainy | 1,048 (40.5) | 774 (29.9) | - | 763 (29.5) | 2,585 (100) |
| Kitambo | Urban | dry | 372 (52.7) | 125 (17.7) | - | 209 (29.6) | 706 (100) |
| Mont-Ngafula | Peri-urban | rainy | 448 (16.9) | 1,416 (53.5) | 2 (0.08) | 781 (29.5) | 2,647 (100) |
| Mont-Ngafula | Peri-urban | dry | 71 (11.1) | 379 (59.3) | - | 189 (29.6) | 639 (100) |
| **Subtotal**<br>**(urban)** |  |  | 1,420 (44.4) | 899 (28.1) | - | 972 (27.5) | 3,291 (100) |
| **Subtotal**<br>**(peri-urban)** |  |  | 519 (15.7) | 1,795 (54.1) | 2 (0.06%) | 970 (29.2) | 3,286 (100) |
| **Total** |  |  | 1,939 (29.5) | 2,694 (41.0) | 2 (0.03%) | 1,942 (29.5) | 6,577 (100) |

### Re-use potential

The dataset from this study provides various entomological data on *Aedes* across multiple area types (urban and peri urban areas), and sampling methods in Kinshasa during rain and dry seasons. It can be directly applied to vector surveillance programmes to identify high risk areas and track mosquito abundance depending on season in both adult and immature stages. The dataset will also support spatial modelling and risk mapping, offering valuable inputs for developing predictive models of *Aedes* mosquitoes species under varying environmental and climatic conditions in in the Democratic Republic of Congo.

## Data Availability

The data supporting this article are published through the IPT of the University of Kinshasa and are available via GBIF under a CC0 waiver [16].

### Dataset description

**Object name:** Darwin Core Archive *Aedes* mosquito distribution across urban and peri-urban areas of Kinshasa city, Democratic Republic of Congo

**Character encoding:** UTF-8

**Format name:** Darwin Core Archive format

**Format version:** 1.0

**Distribution:** https://cloud.gbif.org/africa/archive.do?r=aedes_data

**Publication date of data:** 2025-06-07

**Language:** English

**Licences of use:** Public Domain (CC0 1.0)

**Metadata language:** English

**Date of metadata creation:** 2025-06-07

**Hierarchy level:** Dataset

### Editor’s note

This paper is part of a series of Data Release articles working with GBIF and supported by TDR, the Special Programme for Research and Training in Tropical Diseases hosted at the World Health Organization, in order to publish datasets on vectors of human diseases [20].

## Abbreviations

BIOLAV: Laboratory of Bioecology and Vector Control
GBIF: Global Biodiversity Information Facility
HLC: human landing catches
IPT: Integrated Publishing Toolkit

## Declarations

## Acknowledgements

We thank all participants in this study, in particular the team of the Bioecology and Vector Control Laboratory at the Kinshasa School of Public Health for their technical assistance and support during field and laboratory activities. We acknowledge the commitment of all mosquito collectors and focal points, and we are grateful to the local communities for their collaboration and hospitality during field work. We also thank Tsiky Rabetrano (GBIF Afrique) for his guidance and technical support.

## Ethical approval and consent to participate

Not applicable.

## Authors’ contributions

EM, JZ and VN contributed to the design and coordination of the project, as well as the quality control of the mosquito samples. FA contributed to the coordination for the operational execution of this study in the field. VN and AB contributed to data curation, visualization, formal analysis on excel book, and drafting of the manuscript. VN, NB, NM, GD, DK, LM, HL, AB and CB supervised the field study, conducted laboratory experiments, including mosquito sorting and morphological identifications. All authors read and approved the final manuscript.

